# Behavioral demands organize a decision process into distinct yet coordinated neural representations in parietal cortex

**DOI:** 10.64898/2026.09.11.750920

**Authors:** NaYoung So, Ariel Zylberberg, Michael N Shadlen

**Affiliations:** Zuckerman Mind Brain Behavior Institute, Department of Neuroscience, Columbia University, New York, United States; Howard Hughes Medical Institute; Kavli Institute

## Abstract

Perceptual decisions are widely modeled as the accumulation of evidence to a bound. In the lateral intraparietal area (LIP), this computation is thought to be implemented in a low-dimensional population representation organized around the single action used to report the choice, consistent with an intentional framework. The intentional framework, however, implies that changing the behavioral demands on the report should change the representation itself, raising a question about the generality of the low-dimensional decision representation described in LIP: is it a special case of decisions reported through a single action, or does it reflect a more general computational architecture that can support multiple behavioral outputs? We tested this by training monkeys to report the *termination* of a motion-discrimination decision with a saccade to a choice-neutral target, and its *content* only later, with a saccade to one of two choice targets. Even though the two reports were behaviorally separable, the timing of termination remained systematically linked to the accumulation of sensory evidence supporting the eventual choice in both monkeys, indicating that both reports continued to draw on a common underlying computation. Using high-density Neuropixels recordings from LIP, however, we found that decision termination and decision content were represented along orthogonal population coding directions supported by largely non-overlapping groups of neurons. Yet the two representations were not independent: trial-by-trial fluctuations in the population encoding content predicted subsequent fluctuations in the population encoding termination, with their coupling strengthening as the decision evolved. These results suggest that a single decision computation can be flexibly reformatted into distinct, action-specific representations, coordinated by selective transfer of information between neural populations.

---

Many perceptual decisions are thought to arise from the accumulation of noisy evidence to a bound, at which point the decision terminates and a choice is reported. This drift-diffusion model reconciles the trade-off between decision speed and accuracy, accounting for both the pattern of choices and the distribution of reaction times across a range of task difficulties [1,2,3,4,5]. The underlying neurobiology has been studied extensively using simple, controllable perceptual judgments—most often, discriminating the net direction of a noisy random-dot motion display, whose difficulty is set by the strength of the motion signal [6]. Reported this way, a decision has two facets: content (which direction is the motion?) and termination (when has enough evidence accumulated to decide?)—both conveyed, in the standard task, by a single action, such as an eye movement to one of two choice targets.

In the lateral intraparietal area (LIP) of the macaque, neurons whose response fields overlap the choice target show trial-averaged firing rates that ramp at a rate depending on the strength and direction of the evidence and approach a common level near the time of the response (see [6,7]), consistent with the drift-diffusion process. Because this signature is obtained by averaging across many repetitions of nominally similar trials, however, it reveals only the deterministic component of accumulation; the stochastic component thought to produce variable choices and reaction times has, until recently, been inferred from behavior and from indirect neural signatures rather than observed directly. This association between an evolving decision variable and the activity of neurons tied to a specific action anticipates a broader idea: the intentional framework, which holds that decisions are represented not as abstract, effector-neutral propositions, but as evolving commitments to particular actions—provisional intentions, such as an eye movement to a particular location [8,9].

High-density Neuropixels recordings have begun to close the gap left by trial averaging [10]. Simultaneous recordings from hundreds of LIP neurons resolve, on a single trial, a scalar decision variable that closely approximates drift-diffusion and predicts the choice and reaction time on that very trial [11]. This signal is dominated by a small, identifiable pool of neurons whose response fields overlap the single choice target used to report the decision, providing a direct observation of the single-trial trajectory itself—a different order of evidence from the trial-averaged firing rates and the variance signatures of bounded diffusion previously derived from single-neuron spike counts [12], or from behavioral model-fitting alone.

This single-trial trajectory is low-dimensional, consistent with the intentional framework introduced above. On this view, a low-dimensional code suffices because the decision bears on a single intended report: the same saccade specifies both the content of the decision and the moment it is terminated, so both facets are necessarily carried by the same population of action-specific neurons. In tasks where the duration of the evidence is controlled by the environment rather than the decision-maker, the timing of this covert commitment can be inferred from behavior and dissociated in time from the eventual report [7]; in every such study, however, the report itself remains a single action that simultaneously conveys content and confirms that a decision has been reached.

Low-dimensional structure is observed across many neural systems (e.g., [13,14,15,16]) and has been proposed to facilitate communication and computation through anatomical bottlenecks [17]. The low dimensionality observed in LIP might therefore reflect a general principle of neural information processing. Yet it remains unknown whether, in the context of decision-making, this organization reflects something fundamental about how decisions are computed or instead the structure of the behavioral report. Note that in every study to date, a single action has carried the full weight of the decision. The intentional framework makes a direct prediction here: if dimensionality reflects the number of action-specific pools engaged by a decision, then requiring the decision-maker to report the moment of commitment and the choice with two separate actions should be sufficient to produce an additional, orthogonal dimension.^1^ The observed low-dimensionality of the standard task is, after all, compatible with two very different organizations: a single population that jointly encodes content and termination because they are computed together, or two separable, action-specific populations whose activity has never been dissociated because a single action has always carried both.

Confirming the prediction of the intentional framework would not, by itself, resolve this question. Two distinct population representations, each tied to its own target-specific pool, would be equally compatible with a single evolving decision that is read out onto two action-specific channels, or with two entirely independent processes that happen to terminate in distinct actions. The dimensionality of the neural code and the computational architecture that produced it are therefore separable questions [18,19], and determining how they align may reveal whether, and how, a decision is transformed into distinct behavioral outputs.

Here, we designed a task in which monkeys viewed a random-dot motion stimulus to discriminate its direction and reported the decision termination with a saccade to a choice-neutral target. Only after a set of intervening eye movements did they report the decision itself with a saccade to one of the choice targets. We found that, even when the reports of decision termination and decision content were behaviorally distinct, the timing of termination remained linked to the accumulation of sensory evidence in both monkeys, indicating that the two reports, though separable, arose from a common underlying process.

Using high-density Neuropixels recordings from LIP, we found that decision termination and decision content were represented along orthogonal population coding directions, supported by largely non-overlapping groups of neurons—indicating that the dimensionality of the decision-related code is not fixed, but tracks the structure of the reporting action, consistent with a direct, action-specific code for each report, as the intentional framework predicts. Despite this expected separation, however, trial-by-trial fluctuations in the two representations became increasingly coupled as the decision evolved, and beginning 200–300 ms after motion onset, the representation of decision content increasingly predicted the subsequent representation of decision termination. This temporally asymmetric relationship indicates that, although the two representations drive distinct actions, they remain coordinated through the selective transfer of decision-related information between distinct neural populations. More broadly, these findings suggest that the decision to terminate deliberation and the specification of decision content arise from the same evolving decision process, which can be flexibly reformatted into distinct neural representations, each driving its own action.

## RESULTS

We recorded from 3,566 neurons in the LIP of two rhesus macaques (*Macaca mulatta*) across 31 recording sessions (22 sessions in monkey H, 132.9 ± 5.8 s.e.m. neurons/session; 9 sessions in monkey N, 71.3 ± 14.0 s.e.m. neurons/session) while they performed a motion-discrimination task designed to dissociate the reports of decision timing and decision content (**Figure 1A**). On each trial, the monkey received a liquid reward for correctly identifying the net direction of a random-dot motion (RDM) stimulus displayed at the point of fixation (up vs. down for monkey H; left vs. right for monkey N). However, the monkey was not permitted to report this judgment right away: it first had to make a self-initiated saccade to a choice-neutral blue target, which immediately extinguished the RDM stimulus. This first saccade therefore reported the termination of the decision process without reporting its content, and we refer to the blue target as the *When*-target (*T*^when^) and to the time from motion onset to this saccade as the reaction time (RT). After fixating *T*^when^ and tracking a moving fixation point back to the center of the screen, the monkey then reported the direction of motion with a second saccade to one of two red *What*-targets (*T*^what^). Thus, each trial comprised two sequential saccades: the first reported when the decision terminated, and the second reported what decision was made.

**Figure 1.**
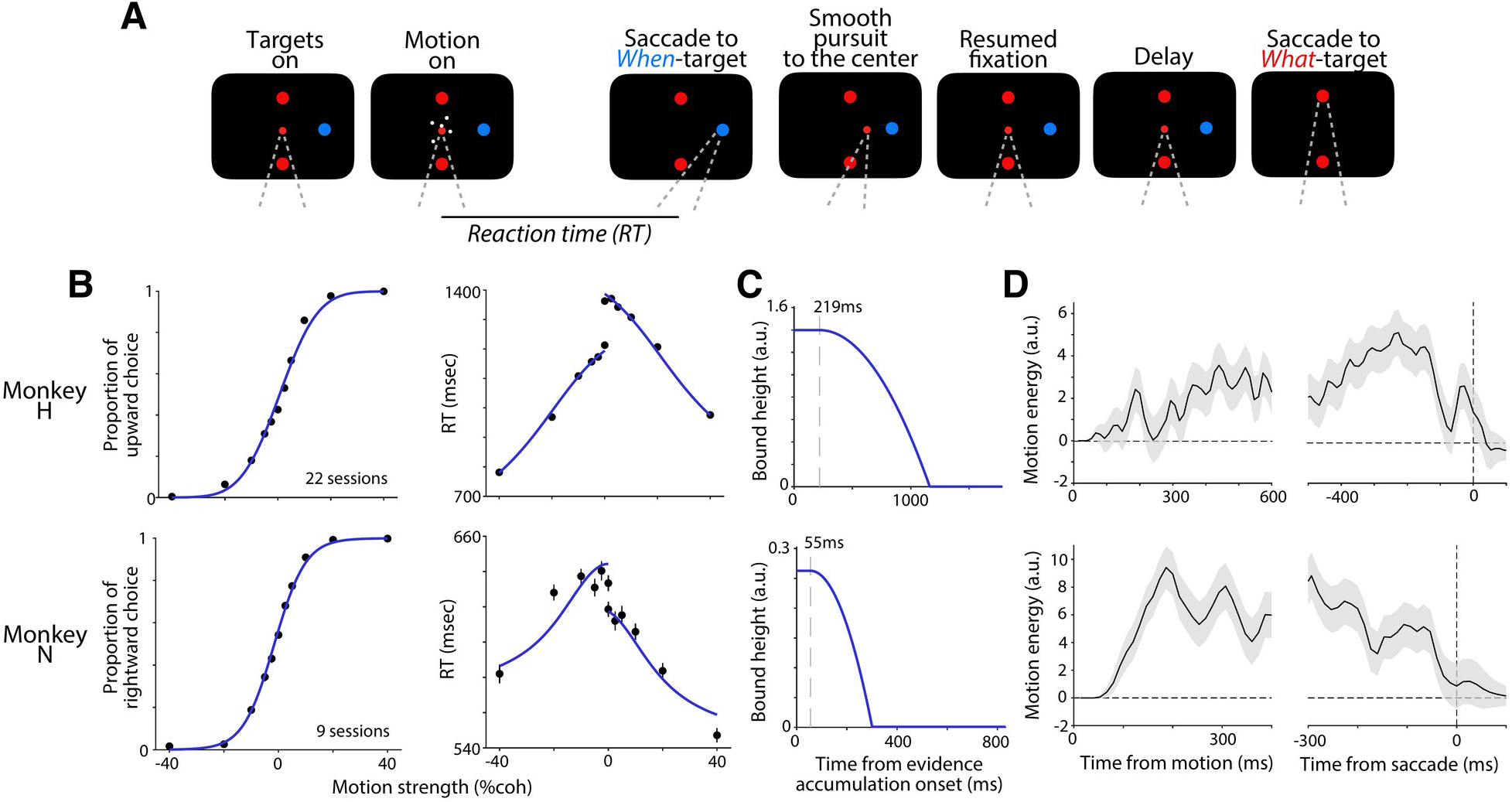
Task and behavior. Monkeys performed a motion-discrimination task in which the timing and content of the decision were reported by separate eye movements. A random-dot motion (RDM) stimulus was presented at the point of fixation (FP) at the center and remained visible until the monkey made a self-initiated saccade to a choice-neutral blue target (*T*^when^), reporting decision termination. After fixating *T*^when^, the monkey made a smooth-pursuit eye movement back to the center, guided by a moving FP. Following a variable delay, the FP was extinguished, and the monkey reported motion direction with a second saccade to one of two red choice targets (*T*^what^), indicating decision content. **A**, Sequence of events in a single trial. The first saccade indicates *when* the decision terminates, and the second saccade indicates *what* decision was made. A key feature of the task is that stimulus viewing duration is self-determined by the monkey, allowing measurement of reaction time (RT; time from motion onset to the saccade) on each trial. The schematic illustrates the task for monkey H (vertical motion discrimination). For monkey N (horizontal discrimination), *T*^when^ was positioned above fixation and the two *T*^what^ were positioned to the left and right. **B**, Behavioral performance. (*Left*) Psychometric functions showing the proportion of choices in the positive direction (up for monkey H; right for monkey N) as a function of signed motion strength. (*Right*) Chronometric functions showing reaction time to the *T*^when^ saccade as a function of motion strength. Curves are fits to a drift diffusion model. Error bars are s.e.; some are smaller than the symbols. **C**, Inferred decision bounds from the model fit for monkey H (*top*) and monkey N (*bottom*). The dashed line indicates the time when the bound begins to decrease for each monkey. **D**, Psychophysical reverse-correlation analysis. Curves show the average motion energy aligned to motion onset (*left*) and first-saccade onset (*right*) for 0% coherence trials, illustrating the influence of moment-to-moment fluctuations in motion evidence on choice. Motion energy is signed positive when consistent with the choice. Shaded regions indicate mean ± s.e.m.

### Separate yet coordinated reports of the When and What of a decision

Choice and RT were both well explained by a single drift-diffusion model, in which noisy motion evidence accumulates to a decision bound and triggers the first saccade, even though the timing of termination and the content of the decision were reported by two separate, non-contemporaneous saccades. Accuracy improved and RT decreased systematically with motion strength, two signatures of the same bounded-accumulation process (**Figure 1B**; lines show model fits). The model included parameters governing the rate of accumulation and decision bounds, as well as additional terms such as time-dependent collapsing bounds, bias parameters, and non-decision time to account for differences in RT distributions across monkeys and choice conditions. **Figure 1C** shows the inferred decision bounds from the best-fitting model, which capture the differences in RT ranges between monkeys.

Despite differences in RT distributions and inferred model parameters between the two monkeys, the behavior of both monkeys indicated that the timing of the initial saccade to *T*^when^ remained closely linked to the same underlying evidence-accumulation process. An alternative model in which decisions terminated according to a predetermined deadline, irrespective of the accumulation process, provided a substantially poorer account of behavior in both monkeys (**Figure S1**). In addition, psychophysical reverse-correlation analyses showed that moment-to-moment fluctuations in motion up to the first saccade influenced choice for both monkeys, supporting temporal integration of evidence (**Figure 1D**).

Despite this link between termination timing and the magnitude of accumulated evidence, termination did not depend on magnitude alone: its timing also depended on which decision was forming. This was particularly evident in monkey H, which showed markedly different RT distributions for the two motion choices. Consistent with this, an alternative model in which termination depended solely on the accumulated magnitude of evidence, irrespective of its direction, failed to account for the joint accuracy and RT distributions in either monkey (**Figure S1**).

Together, these behavioral results show that, although the two saccades reported distinct aspects of the decision—one signaling that a decision had been reached, the other signaling what had been decided—both remained tied to a common, scalar evidence-accumulation process. This raises the question of how this single signal is represented in the brain, and how it gives rise to two separately reported aspects of the decision. We addressed this question with population recordings from LIP.

### Neural representations of the When and What of a decision in LIP

We recorded simultaneously from large populations of LIP neurons—3,566 neurons across 31 sessions (see Methods)—using a high-density Neuropixels probe, an approach inspired by the single-trial resolution of a scalar decision variable achieved with similar recordings in a related task [11]. These neurons had diverse response fields, including many overlapping the *T*^when^ and *T*^what^ target locations. Using population activity during the motion-viewing period, we trained separate decoders to estimate *(i)* whether the first saccade reporting decision termination was imminent at each time point (*When*-decoder) and *(ii)* the monkey’s final choice (*What*-decoder). The resulting weight vectors define coding directions (*When*-CD and *What*-CD, respectively) for each session. Decoders were trained on half of the trials (odd-numbered trials) and evaluated on the remaining trials.

Projection of population activity onto the *When*-CD revealed a ramping pattern, with steeper ramps associated with shorter RTs. **Figure 2A** shows the population activity projected onto the *When*-CD, with trials grouped into six RT bins and averaged across sessions for each monkey. For both monkeys, the slope of the *When* representation was negatively correlated with RT, assessed by pooling the slope estimates and mean RTs for each RT bin across sessions after removing each session’s own mean to account for between-session differences (monkey H: r = -0.75, p = 6.7×10^-25^; monkey N: r = -0.71, p = 5.6×10^-8^). This relationship remained strong when the fastest and slowest RT bins were excluded (monkey H: r = -0.80, p = 1.5×10 ^-20^; monkey N: r = -0.65, p = 1.8×10^-5^), indicating that it is not driven by the extreme RT bins. Notably, in monkey N, the *When* representation exhibited an initial offset that was already correlated with RT prior to motion onset; however, the slope of the ramp remained strongly RT-dependent, indicating that the evolving activity—not merely the pre-motion baseline—reflects the process leading to decision termination.

**Figure 2.**
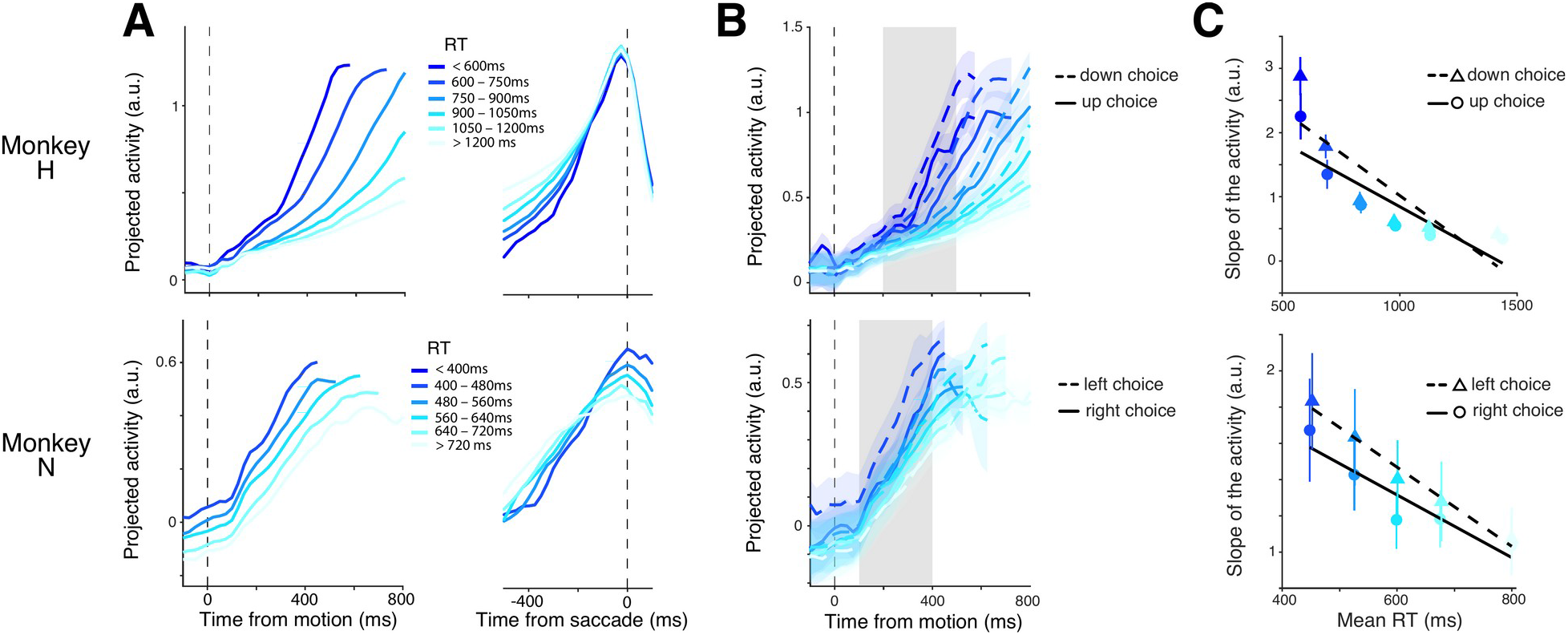
Representation of decision termination in the LIP population. **A**, Population activity was projected onto the *When* coding direction (*When*-CD) to obtain a *When representation* for each trial. Shown are these representations aligned to motion onset (*left*) and the initial saccade to the choice-neutral *T*^when^ (*right*), pooled across sessions, grouped into six RT bins, and averaged within each RT group, for monkey H (*top*) and monkey N (*bottom*). Error trials are included in all panels of Figure 2. **B**, *When* representation, grouped by RT bin (colors) and choice (solid and dashed traces), aligned to motion onset. Traces were averaged within each session for each RT × choice group and then averaged across sessions. Shaded regions indicate mean ± s.e.m. across sessions. Gray shading indicates the time window used to estimate the activity slope. **C**, Slope of the *When* representation during the initial decision epoch plotted against mean RT. Lines show linear regressions relating activity slope to mean RT for each choice condition.

This RT-dependent ramping was preserved when trials were further grouped by choice (**Figure 2B**), indicating that the *When* representation is largely independent of the *What* representation. Consistent with this, the slope of the *When* representation was negatively correlated with RT for both monkeys and for each choice condition (**Figure 2C**), assessed by pooling the slope estimates and mean RTs for each RT bin across sessions after removing each session’s own mean (monkey H: r = -0.67, p = 3.8×10^-16^ for up choices; r = -0.74, p = 2.8×10^-24^ for down choices; monkey N: r = -0.58, p = 2.4×10^-5^ for right choices; r = -0.73, p = 3.0×10^-8^ for left choices). This relationship remained strong when the fastest and slowest RT bins were excluded (monkey H: r = -0.67, p = 1.6×10^-12^ for up choices; r = -0.77, p = 1.4×10^-18^ for down choices; monkey N: r = -0.55, p = 5.0×10^-4^ for right choices; r = -0.69, p = 5.4×10^-6^ for left choices). At the per-session level, the dependence of slope on RT was significantly different from zero for both monkeys and for each choice condition (Wilcoxon signed-rank test: monkey H: p = 4.0×10^-5^ for up choices and p = 4.0×10^-5^ for down choices; monkey N: p = 0.027 for right choices and p = 0.008 for left choices).

The trial-pooled projection onto the *What*-CD revealed a robust representation consistent with an evolving decision variable (DV), which became categorical by the time of the first saccade to *T*^when^, reflecting the monkey’s eventual choice (**Figure 3A**). When aligned to motion onset, this activity differentiated according to both motion direction and strength, with stronger coherence leading to faster separation of the DV; this pattern was preserved when the *What* representation was instead averaged within each session before being averaged across sessions (**Figure 3B**). Consistent with this, the direction-consistent buildup rate was positively correlated with motion strength for both monkeys (**Figure 3C**), assessed by pooling the buildup rate for each motion coherence across sessions after removing each session’s own mean (monkey H: r = 0.61, p = 4.7×10^-15^ for upward motion; r = 0.65, p = 2.7×10^-17^ for downward motion; monkey N: r = 0.32, p = 0.017 for rightward motion; r = 0.82, p = 2.2×10^-14^ for leftward motion), though the effect for rightward motion in monkey N remained markedly weaker than for the other conditions. At the level of individual sessions, the dependence of buildup rate on coherence was significantly greater than zero for both monkeys, with the exception of the rightward-motion condition in monkey N (Wilcoxon signed-rank test: monkey H: p = 5.3×10^-5^ for upward motion and p = 1.0×10^-4^ for downward motion; monkey N: p = 0.164 for rightward motion and p = 0.0039 for leftward motion).

**Figure 3.**
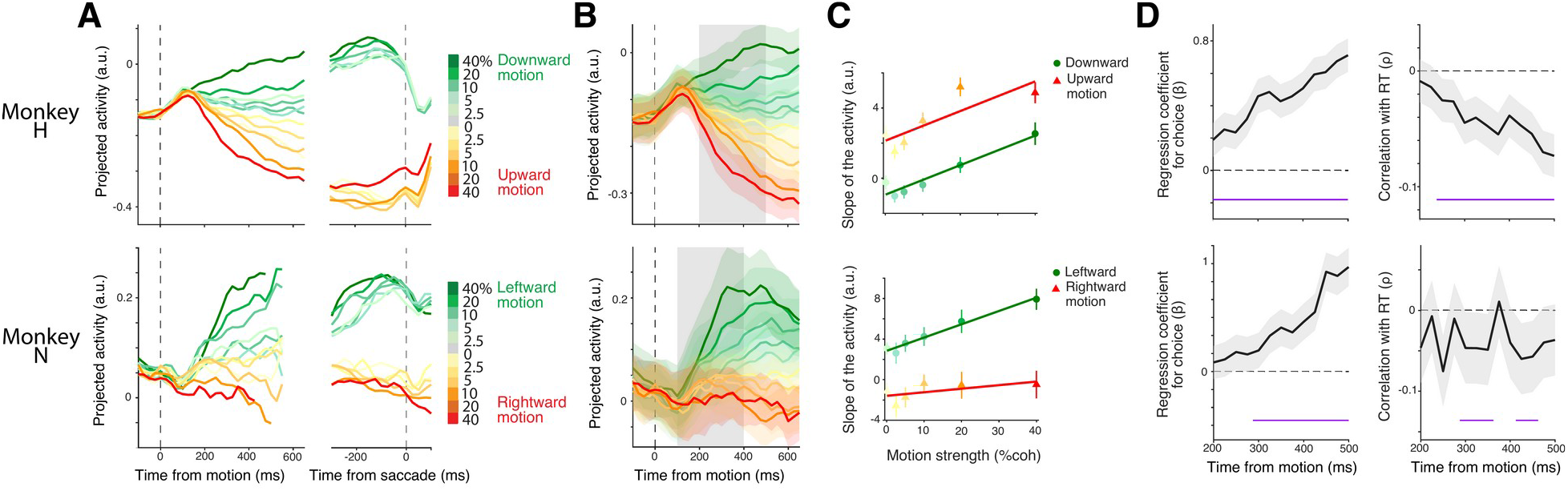
Representation of decision content in the LIP population. **A**, Population activity was projected onto the *What* coding direction (*What*-CD) to obtain a *What representation* for each trial. Shown are these representations aligned to motion onset (*left*) and the initial saccade to the choice-neutral *T*^when^ (*right*), pooled across sessions, grouped by motion direction and strength, and averaged within each motion condition, for monkey H (*top*) and monkey N (*bottom*). Error trials are included throughout Figure 3, except for the panel aligned to the initial saccade ( *right*), where only correct trials are shown to more clearly illustrate the choice-dependent separation of the *What* representation. **B**, *What* representation, grouped by motion direction and strength and aligned to motion onset. Traces were averaged within each session for each motion condition and then averaged across sessions. Shaded regions indicate mean ± s.e.m. across sessions. Gray shading indicates the time window used to estimate the activity slope. **C**, Slope of the *What* representation during the initial decision epoch plotted as a function of motion strength. Lines show linear regressions fit separately for each motion direction. **D**, Relationship between residual *What* representation and behavioral measures over time. Residual activity was computed for each session by subtracting the mean *What* representation for each motion condition, thereby isolating trial-by-trial fluctuations after accounting for stimulus condition. Residualized trials were then pooled across sessions for each monkey. Relationships with the monkey’s eventual choice (*left*; logistic-regression coefficient, β) and RT (*right*; Pearson correlation, ρ) were computed using low-coherence trials (≤ 10%). For the RT analysis, both choices were included for monkey H, whereas only left-choice trials were included for monkey N. Shaded regions indicate 95% confidence intervals. Purple bars indicate time periods significant by cluster-based permutation test (see Methods).

The weaker dependence in the rightward-motion condition for monkey N likely reflects asymmetries in the recorded population, consistent with the placement of the recording chamber in the right hemisphere and the resulting distribution of response fields, which preferentially sampled neurons representing contralateral, leftward choices. In contrast, monkey H performed a vertical motion discrimination task with vertically aligned *T*^what^ targets, likely resulting in a more balanced sampling of neurons representing the two choice alternatives. Hence, for subsequent analyses of the What representation (**Figure 3D** and **Figure 5**), we included only left-choice trials for monkey N, whereas all trials were included for monkey H.

As the decision progressed, trial-by-trial fluctuations in the residual *What* representation increasingly predicted the monkey’s behavior in both monkeys (**Figure 3D**). For each session, residual activity was computed by subtracting the mean *What* representation for each motion condition, isolating trial-by-trial fluctuations after accounting for stimulus condition, and the analysis was restricted to low-coherence trials (≤10%) to further minimize any residual stimulus contribution. The relationship with the monkey’s eventual choice was quantified with a logistic-regression coefficient (β) at each time point, and the relationship with RT was quantified with a Pearson correlation (ρ); time periods of significant association were identified with a cluster-based permutation test (see Methods). The increasing relationship with RT is particularly notable, as it indicates that activity along the *What*-CD progressively reflects information predictive of when the decision will terminate, even though the behavioral reports of *What* and *When* are dissociated.

### Relationship between the When and What coding directions in LIP

The *When*- and *What*-CDs were nearly orthogonal in both monkeys: cosine similarities did not differ significantly from zero (monkey H: mean = 0.014, p = 0.60; monkey N: mean = 0.017, p = 0.79, t-test), and did not differ from a null distribution obtained by randomly permuting the assignment of When-CD weights across neurons within each session—preserving the weights’ distribution and sparsity while disrupting the neuron-by-neuron correspondence between the two coding directions (Wilcoxon signed-rank test comparing each session’s observed cosine similarity to its shuffled mean, 10,000 permutations per session; monkey H: p = 0.69; monkey N: p = 1; **Figure 4**).

**Figure 4.**
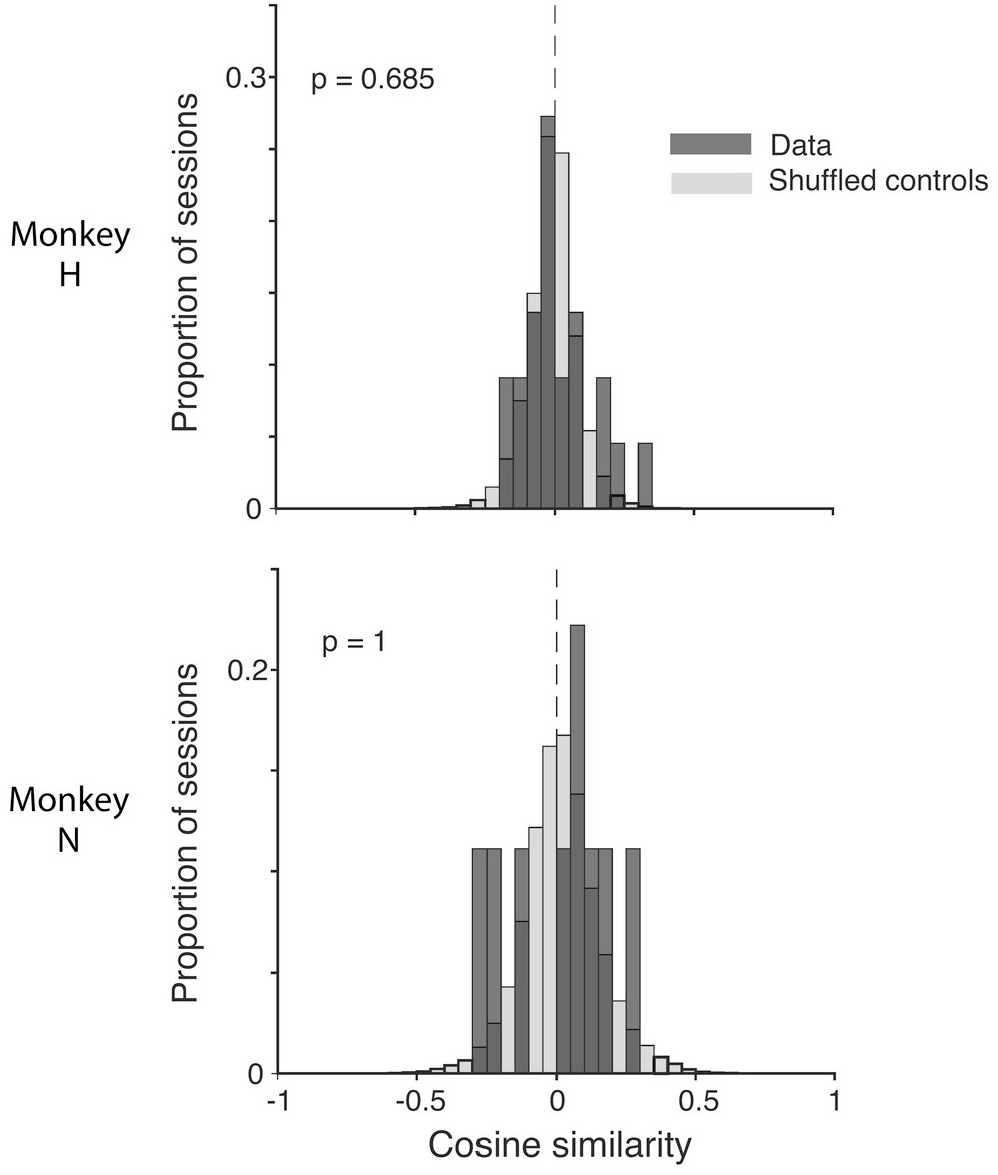
Orthogonal population coding directions for decision termination and decision content. Cosine similarity between the *When* and *What* coding directions (*When*-CD and *What*-CD) computed for each recording session. Dark gray histograms show the distribution of cosine similarities across sessions for monkey H (*top*) and monkey N (*bottom*). Light gray histograms show the null distributions expected from randomly aligned coding directions, obtained by permuting neuron identities 10,000 times within each recording session.

The underlying neuronal populations also showed only minimal overlap. We quantified this overlap using the Jaccard index: the fraction of neurons contributing to either coding direction (i.e., assigned a nonzero decoder weight) that contributed to both. Because neurons contributing to neither CD are excluded from the denominator, this value is greater than or equal to the fraction of the entire recorded population shared between the two CDs. The Jaccard index between neurons with nonzero weights in the two CDs was 0.138 for monkey H and 0.269 for monkey N. This index was not different from a null distribution—obtained by randomly reassigning which neurons contributed to the *When*-CD while holding the *What*-CD’s contributing neurons fixed—for monkey N (Wilcoxon signed-rank test vs. shuffled controls: mean difference = -0.003, p = 0.73), while monkey H showed a slightly higher index than the shuffled controls (mean difference = 0.015, p = 0.02). Restricting to the strongest contributors (top 25% of nonzero weights by magnitude) reduced the overlap further (Jaccard index; monkey H: 0.048; monkey N: 0.053), with no significant deviation from shuffled controls in either monkey (monkey H: mean difference = 0.019, p = 0.08; monkey N: -0.008, p = 1).

This largely orthogonal, minimally overlapping organization is consistent with the prediction from the intentional framework that decision termination and decision content—reported by separate actions—should be carried by largely separate, action-specific populations.

### Interactions between the When and What representations in LIP

The behavioral analyses indicate that decision termination and decision content arise from a common evidence-accumulation process, whereas the neural analyses revealed largely orthogonal population representations for these two aspects of the decision. This is an apparent puzzle: if the two aspects of the decision are carried by largely separate populations, what ties them to the same evolving process? Coordination could arise through concurrent fluctuations driven by a common decision-related signal, in which case the populations representing *When* and *What* would either operate as functionally coupled components of the same decision process or inherit that signal from upstream circuits. Alternatively, coordination could be temporally asymmetric, with activity in one population contributing to the subsequent evolution of the other, consistent with selective transfer of decision-related information between representations.

Trial-by-trial fluctuations in the *What* and *When* representations were positively correlated, but this relationship was temporally asymmetric. In both monkeys, earlier fluctuations in the *What* representation predicted subsequent fluctuations in the *When* representation (**Figure 5A**), consistent with a relationship in which decision-related signals represented along the *What*-CD contribute to the subsequent evolution of the *When* representation. This relationship is unlikely to reflect shared stimulus drive, as the correlation was computed using residual activity obtained by subtracting the mean response expected for each coherence and choice condition within each session.

**Figure 5.**
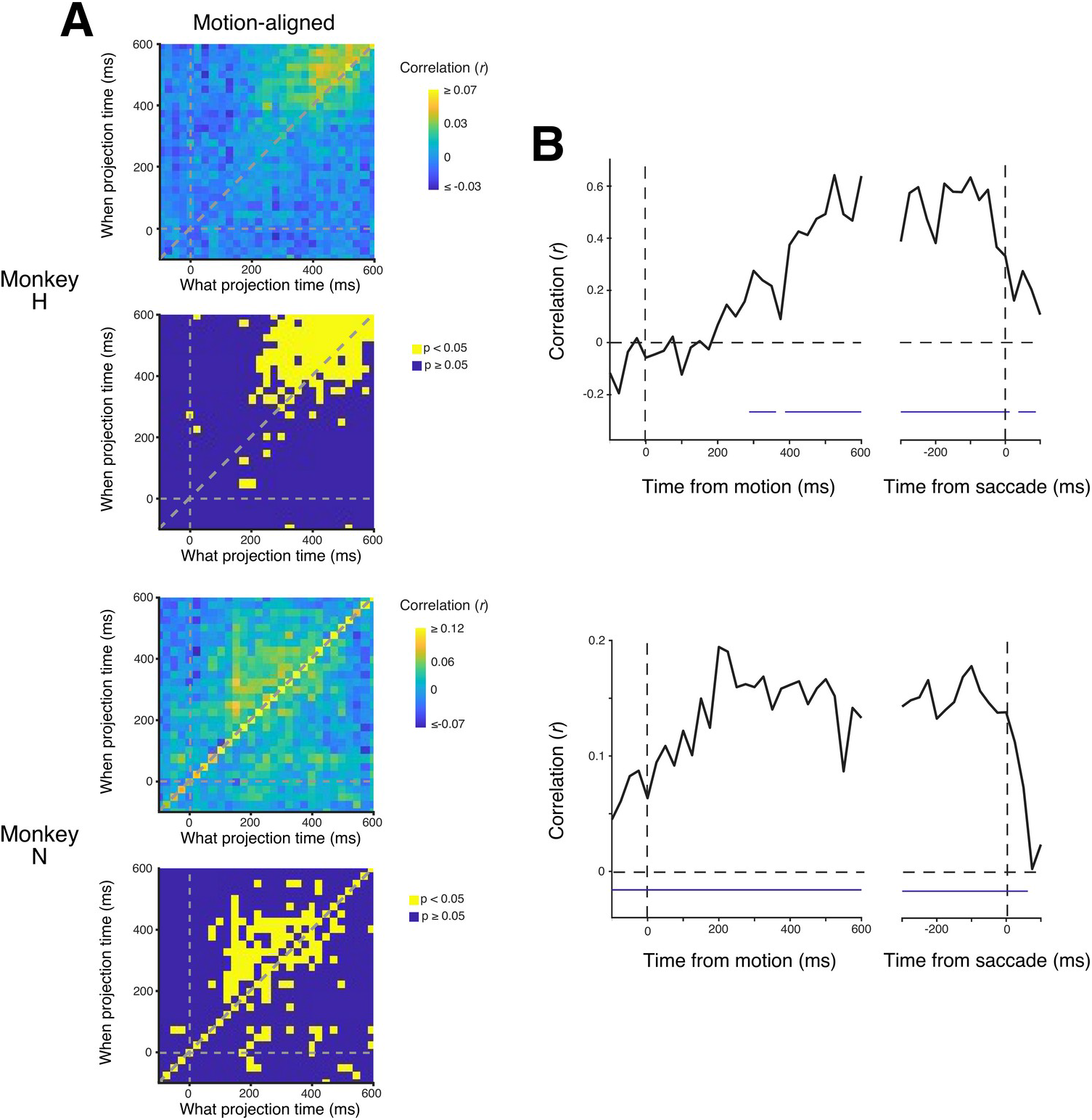
Interactions between the *When* and *What* representations during decision formation. **A**, Trial-by-trial correlation between the residual *What* and *When* representations as a function of time, aligned to motion onset. Residual activity was computed within each session by subtracting the mean representation for each coherence × choice condition. Residual *What* activity was sign-aligned across choices such that positive fluctuations indicated deviations in the direction supporting the monkey’s eventual choice. Residual representations were then pooled across sessions within each monkey, and Pearson correlations were computed across pooled trials for each pair of *What* and *When* time points. All correct-choice trials were included for monkey H, whereas only left-choice correct trials were included for monkey N. Warm colors indicate positive correlations. Lower panels indicate time-point pairs with a positive correlation and p<0.05 (yellow); all other time-point pairs are shown in blue. **B**, Concurrent correlation between the residual *What* and *When* representations (the diagonal of the cross-temporal correlation matrix in **A**) as a function of time, aligned to motion onset (*left*) or the initial saccade to *T*^when^ (*right*). Correlations were computed across trials pooled across sessions within each monkey. Purple bars indicate time periods significant by cluster-based permutation test (see Methods).

The strength of this coupling also evolved systematically over the course of the trial ( **Figure 5B**): the correlation between the two representations increased as the decision progressed and returned toward baseline around the time of the saccade to *T*^when^, when the monkey reported decision termination.

Taken together, these dynamic features indicate a temporally structured interaction between the two representations, consistent with selective transfer of decision-related information between largely distinct populations [20].

## DISCUSSION

We introduced a seemingly minor variation to a highly studied choice-response-time task that has long informed the neurobiology of perceptual decision-making: whereas choice and RT are ordinarily reported through a single action—a saccade to one of two choice targets—our monkeys reported the termination of the decision and its content through two different actions, separated by an enforced delay. Despite this separation, choice and RT remained explained by the same drift-diffusion process that has long accounted for both in the standard, single-action version of the task [21], indicating that termination and content continued to draw on a common evidence-accumulation mechanism even when reported separately. Decision termination and decision content, however, were carried by largely separate populations of LIP neurons. Activity in the population representing decision content, moreover, predicted the subsequent evolution of activity in the population representing decision termination, revealing a temporally asymmetric interaction from the *What* representation to the *When* representation. A single evidence-accumulation process therefore recruits two distinct populations of neurons, one for each action used to report it, with activity in one continuing to shape the other as the decision unfolds.

Although the reports of termination and content were separated in our task, both remained tied to a single evidence-accumulation process, rather than to two independent computations that happen to run on similar timescales. Decision termination, on this view, need not be governed by a process distinct from the one that shapes content; it may instead reflect another readout of the same evolving decision. A similar principle has been proposed in the confidence literature, where confidence and choice are reported separately yet are thought to arise from the same underlying decision variable [22,23]. The same logic extends to content itself: human subjects sometimes reverse an initial choice when evidence that arrives too late to inform the original decision nonetheless arrives in time to revise it before a movement is complete [24]^2^, indicating that a report which appears final at the moment it is registered can still be a readout of the same decision process that continues to evolve. Together, these observations make a broader point: two aspects of a decision can be dissociated behaviorally without reflecting dissociated computations.

Our finding argues against accounts in which termination is governed by an independent timer or deadline process that ends deliberation irrespective of the evolving decision. Our task provides an especially favorable setting for detecting such a mechanism because termination was reported explicitly through an action distinct from the report of decision content. Nevertheless, in both monkeys, the timing of decision termination remained systematically linked to the same evolving decision process that shaped decision content.

Our results are consistent with a prediction of the intentional framework [8,9]: if LIP activity is organized around the actions used to report different aspects of a decision, then even aspects that draw on the same evidence-accumulation process should be represented separately when they are mapped onto distinct actions. This is what we found: decision termination and decision content were represented by largely non-overlapping populations of LIP neurons. Thus, a common decision process can recruit distinct action-specific populations for the different behavioral outputs it must drive.

This conclusion rests on more than the geometric orthogonality of the two coding directions. Data-driven tools such as principal components analysis and choice decoding recover a scalar decision variable as a projection onto a coding direction in a high-dimensional state space, and such projections can satisfy the statistical signatures of accumulation whether the contributing neurons form one interconnected population or several [26]. In the single-saccade version of this task, for example, several such data-driven coding directions render single-trial signals nearly indistinguishable from the one dominated by the identified, target-specific pool, and only the post hoc identification of response fields reveals that a small, functionally defined subpopulation accounts for the signal [11]. A geometric description of this kind does not by itself distinguish a direct code—a quantity carried in the firing rates of an identifiable, functionally defined pool of neurons tied to a specific action—from a distributed code, in which the quantity is recoverable only by applying a set of weights to a heterogeneous population, a distinction connected to a broader debate about direct versus distributed neural representations [26,27,28]. Orthogonality alone would be compatible with a single, shared population of neurons contributing to both representations. The additional finding that the two coding directions draw on largely non-overlapping neurons is what supports the stronger claim that decision termination and decision content are carried by largely distinct populations, rather than merely distinct directions within a shared one.

This view reconciles our findings with the strong case for a low-dimensional decision variable in the standard, single-saccade version of this task, where a signal carried by a small, choice-target-specific pool of neurons accounts for both the deterministic drift and the trial-to-trial diffusion underlying choice and reaction time [11]. A factor-based, state-space characterization of that population—the kind used to identify low-dimensional latent structure in population activity [29]—would report the same low-dimensional organization regardless of whether that structure is carried by a dedicated neuronal pool or distributed across the population; only identifying which neurons contribute distinguishes these possibilities. Rather than treating this low dimensionality as a general property of the decision computation, we take it to reflect the number of action-specific pools recruited to represent a decision, as argued in the Introduction: decomposing a single-action report into two separately reported actions should be sufficient to produce an additional, orthogonal dimension, with no change to the evidence or the underlying computation. A single action recruits a single, action-specific pool; two separately reported actions recruit an additional one. On this view, the resulting increase in dimensionality need not mean that the underlying computation has changed in kind; it requires only a minor adjustment to a direct-coding scheme—in which an additional population is recruited, or enabled, to carry the newly separated component of the decision [20,30]—rather than a shift from a direct to a genuinely distributed code.

A central implication of our study is that carrying decision termination and decision content in separate populations of neurons creates a coordination problem. Although both arose from a common evidence-accumulation process, they were represented by largely distinct populations of neurons in LIP. For behavior to remain coherent, information about the evolving decision must therefore be shared between these populations.

Our results suggest one mechanism by which this coordination may be achieved. Trial-by-trial correlation analyses revealed that activity in the *What* representation preceded activity in the *When* representation and that coupling between the two increased as the decision evolved, supporting a temporally asymmetric interaction in which information carried by the *What* representation contributes to the subsequent evolution of the *When* representation. Such coordination may reflect flexible information transfer between neural populations [20], enabling a common decision process to be reformatted into distinct action-related representations while preserving the information needed to coordinate behavior. This kind of continuous transfer of an evolving decision variable to systems responsible for action is not unique to LIP or to this task: electrically evoked oculomotor commands in monkeys track the accumulating motion evidence well before the decision is reported [31], and stretch-reflex gains in the human arm track the same evolving decision variable during an analogous discrimination task [32]. In both cases, a downstream system responsible for action is continuously informed by the state of an unfinished decision rather than waiting for its completion. The present findings suggest that a related principle operates within a single decision-related area, coordinating two distinct representations rather than a decision area and its motor targets.

### Limitations

Several limitations constrain the interpretation of our findings. The temporal asymmetry between the two representations identifies a directional predictive relationship but does not establish the circuit mechanisms that give rise to it. In addition, both termination and content were expressed through eye movements in our task; whether the same representational organization applies when they are expressed through different effectors remains to be determined. Moreover, our recordings were confined to LIP, whereas decision formation and termination engage a broader network that also includes prefrontal cortex, basal ganglia, and brainstem structures [33,34,35,36,37]. Within this broader network, the superior colliculus (SC) could contribute to the coordination between the *What* and *When* representations described above. A threshold-crossing signal in SC, reflected in a burst of activity that need not itself trigger a saccade [37], could sense that the decision has reached the point of termination and render the population of LIP neurons representing *T*^when^ transiently receptive to a more widely broadcast signal carrying decision content [30]. Such a mechanism could support selective information transfer between the *What* and *When* populations without requiring dedicated, private wiring between them. This account predicts that the timing of the SC threshold-crossing signal should covary, across trials, with the onset of coupling between the *What* and *When* representations. Our trial-pooled cross-correlation analyses establish the temporal ordering of the two signals but cannot resolve this relationship at the single-trial level. Testing this possibility, and determining whether similar principles of information transfer extend beyond a single cortical area, will require simultaneous recordings and causal manipulations across the distributed decision network.

Our findings point to a more general principle: behavioral dissociation does not necessarily imply computational independence. Even when decision termination and decision content are expressed through distinct actions, both remained linked to a common evidence-accumulation process. This observation cautions against interpreting behaviorally distinct variables as independent computations and instead highlights how a common decision process can be reformatted into multiple representations and behavioral outputs. In this view, the critical question is not whether decision termination is represented separately from decision content, but how information derived from a common computation is transformed to support distinct behavioral demands.

## MATERIALS AND METHODS

### Experimental Model and Subject Details

All training, surgery, and recording procedures were in accordance with the National Institutes of Health Guide for the Care and Use of Laboratory Animals (National Research Council, 2011) and approved by the Columbia University Institutional Animal Care and Use Committee. We performed extracellular neural recordings in the area LIP of two adult male rhesus macaques (M. mulatta, 9–10 kg; Monkey H and N, ages 13 and 14). Prior to data collection, the monkeys were fitted with a titanium headpost (Thomas Recordings, Rogue Research) and a PEEK plastic recording chamber (Rogue Research) designed and positioned based on the MRI of each monkey, over area LIP in the left (Monkey H) or the right (Monkey N) hemisphere. These procedures were conducted under general anesthesia in an AALAC-accredited operating facility using sterile techniques and state-of-the-art monitoring.

### Methods Details

The data set comprises 3,566 single neurons from LIP, recorded using a 384-channel Neuropixels 1.0 NHP probe (long, 45 mm shank; IMEC) over 31 sessions (22 sessions in monkey H, 132.9 ± 5.8 s.e.m. neurons/session; 9 sessions in monkey N, 71.3 ± 14.0 s.e.m. neurons/session). For each recording session, the Neuropixels probe was lowered through the guide tube in the grid, advanced to the ventral part of LIP (LIPv; [38]) using a microdrive (Narishige). The experiments were controlled by the Rex system [39] running under the QNX operating system integrated with other devices in real-time. Visual stimuli were displayed on a CRT monitor (Sony GDM-17SE2T, 75 Hz refresh rate, viewing distance 60 cm) controlled by a Macintosh computer running Psychtoolbox [40] under MATLAB (MathWorks). Eye position was monitored by infrared video using an Eyelink1000 system (1 kHz sampling rate; SR Research). Neural data were recorded using SpikeGLX software and synchronized with behavioral data, event codes, and analog eye signals recorded using the Omniplex system (Plexon Inc.).

### Behavioral task

The monkeys are engaged in a random dot motion (RDM) discrimination task and report which direction the random dots are moving (vertical motion for monkey H, horizontal for monkey N). Each trial begins when the monkey fixates a point (FP) at the center of the visual display. After a delay (50–250 ms), three targets appear on the screen (0.5° diameter, 8° eccentricities): one blue target, *T*^when^, for reporting the decision termination, and two red choice targets, *T*^what^. The target configurations differ between monkeys: two vertical locations for *T*^what^ and right *T*^when^ for monkey H, and two horizontal *T*^what^ and up *T*^when^ for monkey N. After a variable delay from the target onset, a dynamic RDM stimulus is presented at the point of fixation. The RDM comprises a sequence of video frames of random dots (2×2 pixels) within an invisible aperture (5° diameter) to achieve an average dot density of 16.7 dots/deg^2^/s. Motion strength (coherence) is defined as the probability that a dot plotted in one frame is displaced consistent with a fixed velocity (5°/s) toward the direction of net motion; coherence takes signed values, where positive values indicate upward or rightward motion (for monkey H and monkey N, respectively).

The RDM stimulus remains visible while the monkey maintains fixation and is extinguished immediately upon the monkey’s self-initiated saccade to the choice-neutral blue target, *T*^when^. This first saccade reports the termination of the decision process, without reporting its content. Reaction time (RT) is defined as the time from motion onset to the initiation of this saccade.^3^ After fixating *T*^when^, the monkey makes a smooth-pursuit eye movement back to the center of the screen, guided by a moving fixation point. This intervening pursuit epoch introduces a temporal separation between the reports of decision termination and decision content, without requiring prolonged fixation at *T*^when^, and ensuring that both reports are initiated from central fixation. This recentering preserved the geometry of the choice-report saccades, such that the saccade vector associated with each *T*^what^ was the same during motion viewing and at the time of the eventual choice report. Following a variable delay, the FP is extinguished, cueing the monkey to report the perceived direction of motion with a second saccade to one of two red choice targets, *T*^what^. Correct choices are rewarded with juice; on 0% coherence trials, reward is delivered on a random half of trials.

### Analysis of behavioral data

#### Fitting a bounded evidence accumulation model to behavior

We modeled decision formation as a bounded drift-diffusion process in which noisy momentary evidence with mean v = κ C+b (C, signed motion coherence; κ, drift scale; b, a fixed bias) and unit variance accumulated until reaching one of two decision bounds. To allow for urgency-like changes in the bound over the course of a trial, the bounds collapse from an initial height B toward zero: ±[B - B_β_max(t-B_α_,0)^2^] (floored at a small positive value), where B_α_ is the time at which the bound begins to collapse and B_β_ sets the rate of collapse. The saccade to *T*^when^ is triggered by the first passage of the accumulated evidence to either bound; *T*^what^ (the sign of the terminating bound) is reported by the second saccade, after the intervening smooth-pursuit return to the original fixation (Behavioral task). Choice probabilities and choice-conditional mean decision times were obtained by numerically solving the associated Fokker-Planck equation using a spectral method [5]. Predicted RT (the interval from motion onset to the *T*^when^ saccade) is the sum of the modeled decision time and a non-decision time, t_nd_ ^+^ or t_nd_ ^-^, fit separately for trials terminating at each bound (T^+^/T^-^, corresponding to which *T*^what^ target is subsequently chosen).

Model parameters (κ, b, B, B_α_, B_β_, t_nd_ ^+^ and t_nd_ ^-^) were fit by minimizing a joint cost comprising a Gaussian negative log-likelihood on the mean RT for T^+^ and T^-^ trials at each coherence (weighted by the squared s.e.m.) and a binomial negative log-likelihood on the observed choice proportions, using bounded simplex optimization (`fminsearchbnd`). Fits were performed separately for each monkey, pooling trials across all recording sessions.

#### Model comparison

To test whether the timing of the *T*^when^ saccade depends on the accumulated evidence, as opposed to being determined independently of it, we compared the bounded-accumulation model above against two alternatives, fit with the same joint cost function so that all three models’ likelihoods are directly comparable.

##### Imposed deadline

This model retains the same evidence accumulator (drift v=κ C+b) but removes the bound: evidence accumulates freely, and the *T*^when^ saccade is triggered at a stopping time T drawn from a Gamma distribution T Gamma(μ_T_,σ_T_), independently of C. *T*^what^ is reported as the sign of the accumulated evidence at T. This model instantiates the null hypothesis that termination timing does not depend on the evidence at all.

##### Parallel, motion strength-driven timing

This model decouples *T*^when^ from *T*^what^ entirely: a separate “when” accumulator, driven by unsigned motion strength (drift=κ_time_(|C|+ε), ε a small fixed floor) with its own collapsing bound (B_time_, B_α,time_, B_β,time_), determines the time T of the *T*^when^ saccade via its first passage to bound. An independent “what” accumulator (drift v=κ_choice_C+b) is never bounded and is simply read out (its sign taken) at T to determine *T*^what^. This model allows termination time to depend on motion strength—and thus to produce coherence-dependent RT—without termination depending on the signed evidence that determines choice.

In both alternative models, the two non-decision times (t_nd_^+^, t_nd_^-^) were constrained to be equal (**Figure S1**), so that neither model carries any choice-dependent asymmetry—the strictest, most conservative instantiations of “termination independent of evidence,” since allowing these models a choice-dependent (but coherence-independent) non-decision time could let them partially mimic the choice-dependent RT difference that is the diagnostic signature of the coupled model above. As a robustness check, we additionally re-fit both alternative models with separate non-decision times for each choice, matching the coupled model’s flexibility exactly, to confirm our conclusions did not depend on that model alone having this additional degree of freedom (**Figure S2**).

Because all three models share the same cost function, we compared them using both the Akaike [42] and Bayesian Information Criteria [43], AIC=2k+2 NLL and BIC=kln(n)+2 NLL, where k is the number of free parameters (5, 7, and 7 for the deadline, parallel-timing, and coupled models, respectively, in the primary comparison; 6, 8, and 7 in the matched-flexibility robustness check) and n is the number of psychometric/chronometric data points entering the joint cost (the number of coherence levels with a valid choice proportion, plus the number of coherence levels with a valid mean RT for each of the two choices).

#### Psychophysical reverse correlation

To ask whether moment-to-moment fluctuations in the stochastic motion stimulus influenced the monkey’s report, independent of any systematic signal, we analyzed trials at zero coherence (C=0), during which there’s no net motion.

For each trial, motion energy was computed by convolving the actual, frame-by-frame sequence of displayed dot positions with a set of space-time-separable filters implementing the motion energy model of [44]. The dot positions on each frame were rendered as a binary image (each dot a 2×2-pixel square) and rotated so that the trial’s nominal direction of motion was aligned with the filter bank’s x-axis. Spatial filters were an even/odd quadrature pair of 4th-order Cauchy functions (σ=0.5°) along the direction of motion, weighted by an orthogonal Gaussian envelope (σ=0.08°); temporal filters were a “fast” and “slow” pair of impulse-response functions (difference-of-Poisson form, orders 3 and 5, shared time constant k=60 s^-1^; [7,44]). Thirty blank frames—at least as long as the temporal filters’ impulse response—were appended before and after each trial’s stimulus movie prior to filtering, then trimmed from the output. Combining the spatial and temporal quadrature pairs yielded two filters tuned to rightward and two tuned to leftward motion; convolution with the stimulus movie was performed in the frequency domain for computational efficiency. Squaring and summing each direction’s quadrature-pair responses removed the dependence on the exact spatial phase of the dots, giving right- and left-motion energy at every pixel and frame; subtracting (right minus left) and summing over space yielded a single momentary motion energy value per video frame, sampled at the 75 Hz display refresh rate. Because motion energy was computed from the literal dot positions shown on each trial rather than from the trial’s nominal coherence, it captures the stochastic, trial-by-trial fluctuations in net motion around that nominal value.

For each trial, the sign of the motion energy was adjusted according to the monkey’s eventual choice, such that positive motion energy indicated momentary evidence consistent with that choice and negative motion energy indicated evidence inconsistent with it. The resulting motion-energy traces were averaged across trials, both aligned to motion onset and to the initial saccade to *T*^when^. Shaded regions indicate mean ± s.e.m. across trials at each time point.

### Analysis of neural data

Single neurons were identified using Kilosort 4.0 and manually curated in Phy. We additionally excluded neurons whose firing rate did not exceed 6 Hz on at least 50 trials during the motion-viewing epoch, the period on which our analyses focus, to restrict analyses to task-relevant neurons. After these criteria, our main data set comprised 3,566 neurons over 31 recording sessions (22 sessions in monkey H, 132.9 ± 5.8 s.e.m. neurons/session; 9 sessions in monkey N, 71.3 ± 14.0 s.e.m. neurons/session.) Isolation quality was further verified by visual inspection of each unit’s PSTH.

Our main neural analyses consisted of three steps. First, we identified population coding directions associated with decision termination (*When*) and decision content (*What*). Second, we projected population activity onto these directions to obtain continuous neural representations. Third, we quantified how the dynamics of these representations related to decision behavior.

#### Identification of the When CD

To identify the population activity associated with impending decision termination, we trained a decoder to estimate whether the first saccade to *T*^when^ would occur within the next 150 ms. For each recording session, spike counts of all simultaneously recorded neurons were binned in 25 ms bins from 200 ms before motion onset up to 50 ms before the first saccade to *T*^when^ on each trial. Each time bin from each trial was labeled 1 if it fell within 150 ms of that trial’s first saccade to *T*^when^ and 0 otherwise. A single time-independent decoder was then fit, pooling all labeled bins across time and trials within a session, using L1-regularized (LASSO) logistic regression (`lassoglm`, binomial link, fixed regularization λ=0.01). L1 regularization encouraged sparse decoder weights and facilitated interpretation of the resulting coding direction [45]. Only trials with RT exceeding a monkey-specific cutoff (550 ms for monkey H, 400 ms for monkey N) were included. The decoder was fit on a held-out half of trials (a fixed odd/even trial split) and its performance evaluated on the complementary half.

The resulting vector of neuron weights defines the *When* coding direction (*When* CD): the population activity at any time point, projected onto this vector (population state cdot weight vector), gives a continuous scalar readout of how close the population is to the state that precedes the saccade to *T*^when^. This projection was computed at 25 ms resolution.

#### Estimation of the buildup rate of When representation

To quantify how the *When* representation relates to decision timing, we sorted trials into six RT groups using monkey-specific RT bin boundaries ([0 600 750 900 1050 1200 2000] ms for monkey H; [0 400 480 560 640 720 2000] ms for monkey N). For the visualization in **Figure 2A**, single-trial *When* representations were pooled across recording sessions, and the mean representation was computed by averaging across all pooled trials for each RT group.

To assess the relationship between buildup rate and RT, we instead estimated the buildup rate separately for each recording session and RT group, fitting an ordinary least-squares line over a 300 ms window (200–500 ms from motion onset for monkey H; 100–400 ms for monkey N). These session-level buildup-rate and RT estimates were pooled across sessions and RT groups after subtracting each session’s own mean from both quantities, and the Pearson correlation between them was computed separately for each monkey; this demeaning step ensured that the correlation reflected the relationship within sessions rather than being inflated by differences between sessions. To confirm the correlation was not driven solely by the fastest and slowest RT groups, we repeated it after excluding them.

To determine whether the RT-dependent buildup of the *When* representation was independent of decision content, we repeated this analysis after additionally grouping trials by choice. For the visualization in **Figure 2B**, the mean traces were estimated separately for each recording session and RT × choice group, and then averaged across sessions (mean ± s.e.m.). Correspondingly, buildup rates were estimated for each session and RT × choice group separately. For **Figure 2C**, these session-level buildup rates were averaged across sessions for each RT × choice group and plotted against the corresponding mean RTs, together with a linear regression fit to these mean points for each choice condition. Because this group-mean regression relates only six points per choice condition, it can be strongly influenced by any single group mean. To obtain a more robust correlation statistic, we additionally pooled the session-level buildup-rate and RT estimates after subtracting each session’s own mean from both quantities, and computed the Pearson correlation between them separately for each monkey and choice condition; as a control, we also repeated this correlation after excluding the fastest and slowest RT groups.

Finally, to determine whether the relationship between buildup rate and RT was also evident at the level of individual recording sessions, we fit a linear regression relating buildup rate to group-mean RT within each session and choice condition, yielding one regression slope per session. These session-level slopes were tested against zero across sessions using a Wilcoxon signed-rank test.

#### Identification of the What CD

For each recording session, we trained a separate L1-regularized logistic decoder of choice in 100 ms windows stepped every 50 ms, aligned either to motion onset (200–600 ms after motion onset) or to the *T*^when^ saccade (-500 to 0 ms). Training, cross-validation, and trial-inclusion criteria were identical to those used for the *When* decoder. This yielded a family of *What* decoders for each session. From this family, we selected a single *What* decoder for each monkey that generalized broadly across the decision epoch while being sufficiently separated from the *T*^when^ saccade to minimize contributions from peri-saccadic activity (monkey H: 300 ms before the *T*^when^ saccade; monkey N: 500 ms after motion onset). Temporal generalization was assessed by evaluating each decoder on population activity at all time points (**Figure S3**). The weight vector of the selected decoder defined the *What* CD, and projections of population activity onto this vector yielded the *What* representation used in all subsequent analyses.

#### Estimation of the buildup rate of the What representation

To quantify how the *What* representation relates to the strength of sensory evidence, we grouped trials by motion direction and coherence. For the visualization in **Figure 3A**, single-trial *What* representations were pooled across recording sessions, and the mean representation for each motion condition was computed by averaging across all pooled trials.

For the visualization in **Figure 3B**, the mean traces were estimated separately for each recording session and motion condition, and then averaged across sessions (mean ± s.e.m.). To assess the relationship between buildup rate and motion strength, we estimated the buildup rate separately for each recording session and motion condition, fitting an ordinary least-squares line over a 300 ms window (200–500 ms from motion onset for monkey H; 100–400 ms for monkey N).

To compare buildup rates across opposite motion directions, the sign of each session-level buildup-rate estimate was inverted for one direction (upward motion for monkey H; rightward motion for monkey N), so that positive values corresponded to increasing evidence accumulation supporting the motion direction on screen. For **Figure 3C**, these sign-adjusted, session-level buildup rates were averaged across sessions and plotted against motion strength, together with a linear regression fit to these mean points for each motion direction. Because this group-mean regression relates only six points per motion direction, it can be strongly influenced by any single group mean. To obtain a more robust correlation statistic, we additionally pooled the session-level, sign-adjusted buildup-rate and motion-strength estimates across sessions and coherence levels after subtracting each session’s own mean from both quantities, and computed the Pearson correlation between them separately for each monkey and motion direction; to confirm the correlation was not driven solely by the weakest and strongest coherence levels, we repeated it after excluding them.

Finally, to determine whether the relationship between buildup rate and motion strength was also evident at the level of individual recording sessions, we fit a linear regression relating sign-adjusted buildup rate to motion strength within each session and motion direction, yielding one regression slope per session. These session-level slopes were tested against zero across sessions using a Wilcoxon signed-rank test.

#### Relationship between trial-by-trial fluctuations in the What representation and behavior

To determine whether trial-by-trial fluctuations in the *What* representation were related to decision content and timing, we examined their relationship with the monkey’s eventual choice and RT. Analyses were restricted to trials with low motion coherence (≤ 10%) and RT above the monkey-specific cutoff (650 ms for monkey H; 550 ms for monkey N), to minimize stimulus-driven differences and ensure that the analysis window was sufficiently separated from the saccade to *T*^when^.

For each recording session, we computed residual *What* activity by subtracting, at each time point, the corresponding mean *What* representation for each motion condition (direction and coherence). This residualization removed the mean stimulus-dependent component of the *What* representation, allowing us to assess whether remaining trial-by-trial fluctuations were predictive of behavior beyond that expected from the stimulus. Residualized trials were then pooled across recording sessions separately for each monkey.

To quantify the relationship with decision content, we fit a binomial logistic regression predicting the monkey’s eventual choice from the residual *What* representation separately at each time point. The resulting regression coefficient, β, was used as a signed measure of the relationship between trial-by-trial fluctuations in the *What* representation and choice (**Figure 3D**, *left*). The 95% confidence interval for β was computed as β ± 1.96 SE.

To quantify the relationship with decision timing, we first residualized RT within each motion condition by subtracting the mean RT of the corresponding condition. Because the sign of the *What* representation reverses with motion direction, residual *What* activity was sign-aligned across the two motion directions before analysis, such that fluctuations in the decision-consistent direction had the same sign. We then computed the Pearson correlation between the sign-aligned residual *What* representation and residual RT across pooled trials at each time point (**Figure 3D**, *right*). Both choices were included for monkey H, whereas the analysis for monkey N was restricted to left-choice trials. The 95% confidence interval for the correlation coefficient, ρ, was obtained using the Fisher z transformation.

For both analyses, statistical significance over time was assessed using cluster-based permutation tests [46]. Behavioral labels (choice or residual RT) were randomly permuted across trials using the same permutation at all time points, thereby disrupting the trial-by-trial relationship between neural activity and behavior while preserving the temporal structure of neural activity within each trial. Candidate clusters were defined as contiguous time points exceeding a two-sided pointwise threshold of p<0.05, and cluster mass was calculated as the sum of the absolute test statistics within each cluster. Statistical significance was determined from the null distribution of the maximum cluster mass across permutations (5,000 permutations for RT; 1,000 for choice). Fewer permutations were used for the choice analysis because each permutation required refitting the logistic regression at every time point.

#### Relationship between When and What coding directions

To characterize the relationship between the population codes for decision termination and decision content, we compared the *When*- and *What*-CDs within each recording session. Coding directions were represented by the normalized decoder-weight vectors described above. We first quantified their geometric relationship using cosine similarity, which ranges from -1 to 1, with -1 indicating perfectly anti-aligned coding directions, 0 indicating orthogonal directions, and 1 indicating perfectly aligned directions. Cosine similarities were computed separately for each recording session and tested against zero across sessions using a one-sample t-test.

To determine whether the observed alignment differed from that expected by chance, we generated a session-specific null distribution by randomly permuting the assignment of *When*-CD weights across neurons while leaving the *What*-CD weights unchanged. This procedure preserved the distribution and sparsity of the decoder weights while disrupting the neuron-by-neuron correspondence between the two coding directions. The permutation was repeated 10,000 times for each session. The observed cosine similarity for each session was compared with the mean cosine similarity of its shuffled distribution using a Wilcoxon signed-rank test across sessions.

We next quantified the extent to which the same neurons contributed to the *When*- and *What*-CDs. Because L1 regularization produced sparse decoder-weight vectors, we first defined contributing neurons as those assigned nonzero weights and quantified the overlap between the sets of contributing neurons for the two CDs. We additionally assessed overlap among the strongest contributors by restricting each set to the top 25% of its nonzero weights, ranked by absolute weight magnitude. Overlap was quantified using the Jaccard index,

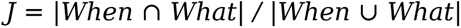

where When and What denote the sets of contributing neurons for the *When*- and *What*-CDs, respectively. For each overlap analysis, chance levels were estimated by randomly reassigning the identities of the contributing *When* neurons within each session while preserving their number and holding the set of contributing *What* neurons fixed. The Jaccard index was recomputed for 10,000 randomizations, yielding a session-specific null distribution that preserved the sizes of the contributing populations. The observed overlap for each session was compared with the mean of its shuffled distribution using a Wilcoxon signed-rank test across sessions. Analyses were performed separately for each monkey.

#### Relationship between When and What representations

To characterize the temporal relationship between the *When* and *What* representations during decision formation, we examined trial-by-trial correlations between residual activity projected onto the two coding directions. Residual representations were computed separately within each recording session. For each coherence × choice condition, we subtracted the corresponding mean *When* and *What* trajectories from the individual-trial trajectories at each time point, thereby removing condition-dependent mean activity while preserving trial-by-trial fluctuations. All correct-choice trials were included for monkey H, whereas only left-choice correct trials were included for monkey N.

Because the sign of the *What* representation depends on the eventual choice, residual *What* activity was sign-aligned across choices such that positive fluctuations indicated deviations in the direction supporting the monkey’s eventual choice. Residual *When* activity was not sign-aligned. Following residualization and sign alignment, single-trial residual representations were pooled across recording sessions separately for each monkey.

We then computed Pearson correlations across the pooled trials between the residual *What* representation at every time point and the residual *When* representation at every time point, yielding a two-dimensional cross-temporal correlation matrix for activity aligned to motion onset (**Figure 5A**). Each element of the matrix quantifies the relationship between trial-by-trial fluctuations in the *What* representation at one time point and fluctuations in the *When* representation at another time point. The corresponding p value for each time-point pair was obtained from the Pearson correlation test.

To characterize the evolution of concurrent coupling, we extracted the diagonal of the motion-onset-aligned cross-temporal correlation matrix, corresponding to correlations between the *When* and *What* representations evaluated at the same time point. We additionally computed this concurrent correlation directly for activity aligned to the initial saccade to *T*^when^ (**Figure 5B**).

To assess the statistical significance of the concurrent correlation over time while accounting for multiple comparisons across adjacent time points, we used a cluster-based permutation test. For each permutation, the correspondence between residual *When* and *What* trials was randomly shuffled while preserving each trial’s complete time course, thereby disrupting trial-by-trial coupling between the two representations while preserving their temporal autocorrelation. Pearson correlations and the corresponding t statistics were recomputed at each time point. Contiguous time points exceeding a two-sided cluster-forming threshold were grouped into clusters, and cluster mass was defined as the sum of the absolute t statistics within each cluster. The maximum cluster mass from each of 5,000 permutations formed the null distribution against which clusters in the observed data were evaluated.

## ACKNOWLEDGEMENTS

We thank Brian Madeira, Cornel Duhaney, Amanda Rodriguez for technical support and animal care. We thank members of Shadlen lab for critical input throughout the research.

## AUTHOR CONTRIBUTIONS

N.S. conceived and designed the experiment and collected the data. N.S. and A.Z. analyzed the data. N.S. and M.N.S. wrote the initial draft and all authors edited the manuscript. M.N.S. supervised the project.

## DATA AND CODE AVAILABILITY

Data and code supporting the findings of this study will be made available upon publication.

## SUPPORTING INFORMATION

### Supplementary figures

**Figure S1.**
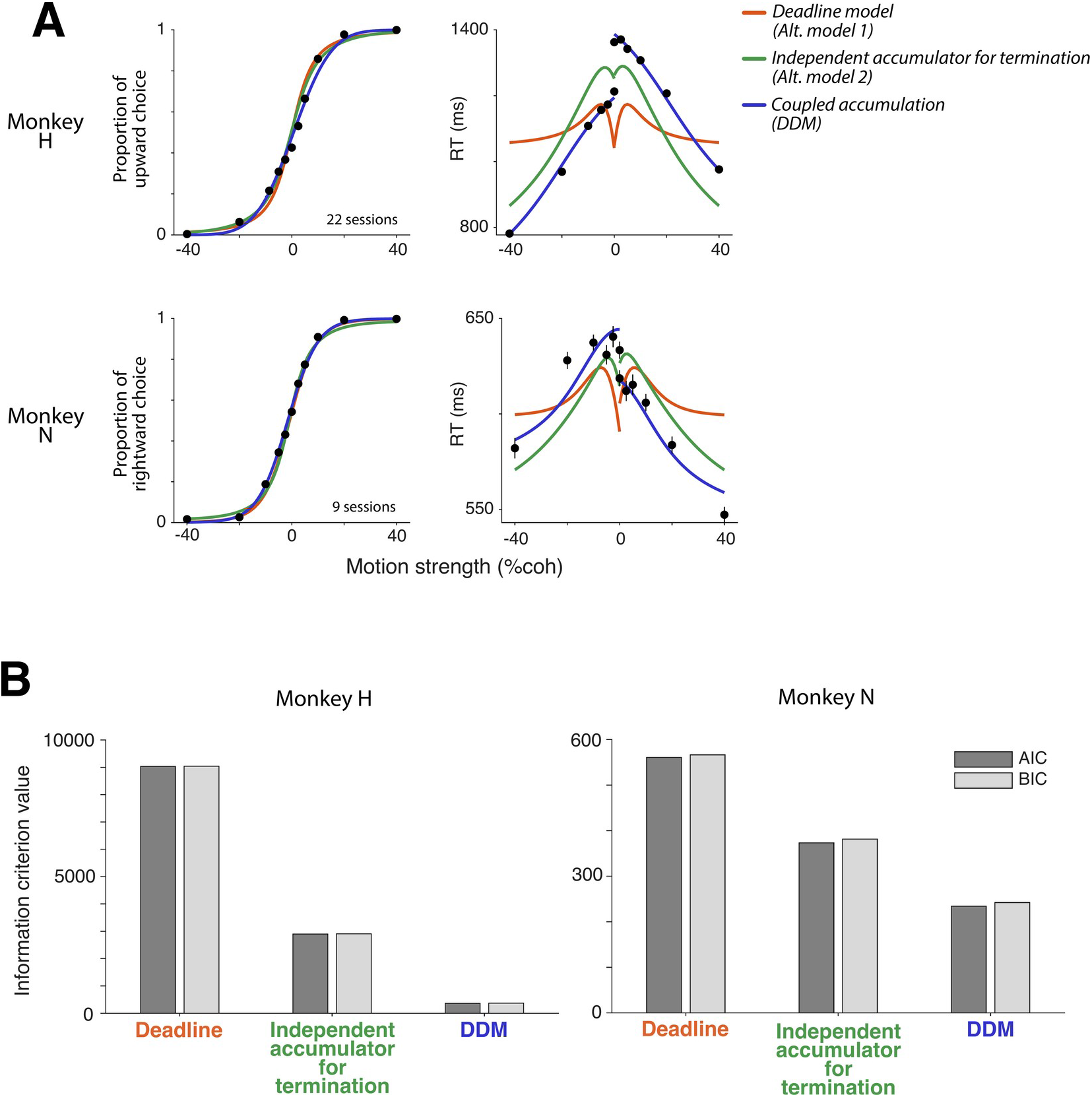
Comparison of coupled and independent-termination models. **A,** Psychometric functions (*left*) and choice-conditional chronometric functions (*right*) for monkey H (*top*) and monkey N (*bottom*), together with fits of three models of decision termination and choice: the *Deadline* (Alt. model 1), *Independent accumulator for termination* (Alt. model 2), and *Coupled accumulation* (DDM) models. In the *Deadline* model, evidence accumulates without a bound and termination occurs at an independently generated stopping time. In the *Independent accumulator for termination* model, termination is determined by a separate accumulator driven by unsigned motion strength, while choice is determined by an independent signed-evidence accumulator evaluated at the time of termination. In the *Coupled accumulation* model, signed evidence accumulates to one of two collapsing bounds, with bound crossing jointly determining the time of decision termination and the subsequently reported choice. For the two alternative models, non-decision time was constrained to be the same for the two choices (t_nd_^+^=t_nd_^-^); Figure S2 relaxes this constraint. **B,** Akaike information criterion (AIC) and Bayesian information criterion (BIC) for the three models for monkey H (*left*) and monkey N (*right*). Lower values indicate better model performance after accounting for model complexity.

**Figure S2.**
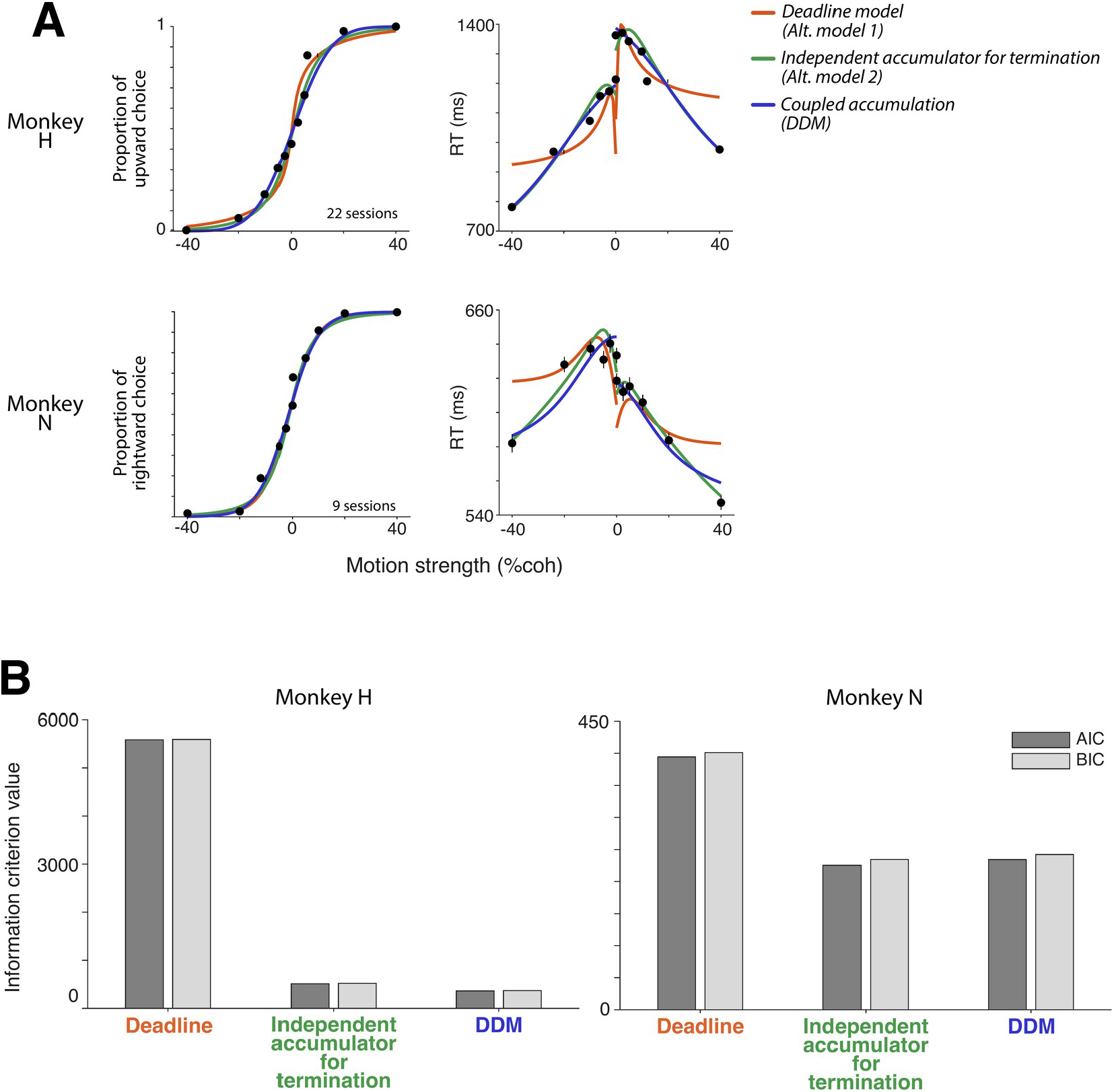
Model comparison allowing choice-dependent non-decision times in the alternative models. Same model architectures and conventions as in Figure S1 except that the *Deadline* (Alt. model 1) and *Independent accumulator for termination* (Alt. model 2) models were allowed separate non-decision times for the two choices (t_nd_^+^ and t_nd_^-^), matching the flexibility of the *Coupled accumulation* (DDM) model. **A,** Psychometric functions (*left*) and choice-conditional chronometric functions (*right*) for monkey H (*top*) and monkey N (*bottom*), together with fits of the three models. Allowing separate non-decision times provides the alternative models with an additional source of choice-dependent RT differences. **B,** AIC and BIC comparisons for the three models for monkey H (*left*) and monkey N (*right*). With this additional flexibility, the *Independent accumulator for termination* model was modestly preferred for monkey N, whereas the *Coupled accumulation* model remained preferred for monkey H.

**Figure S3.**
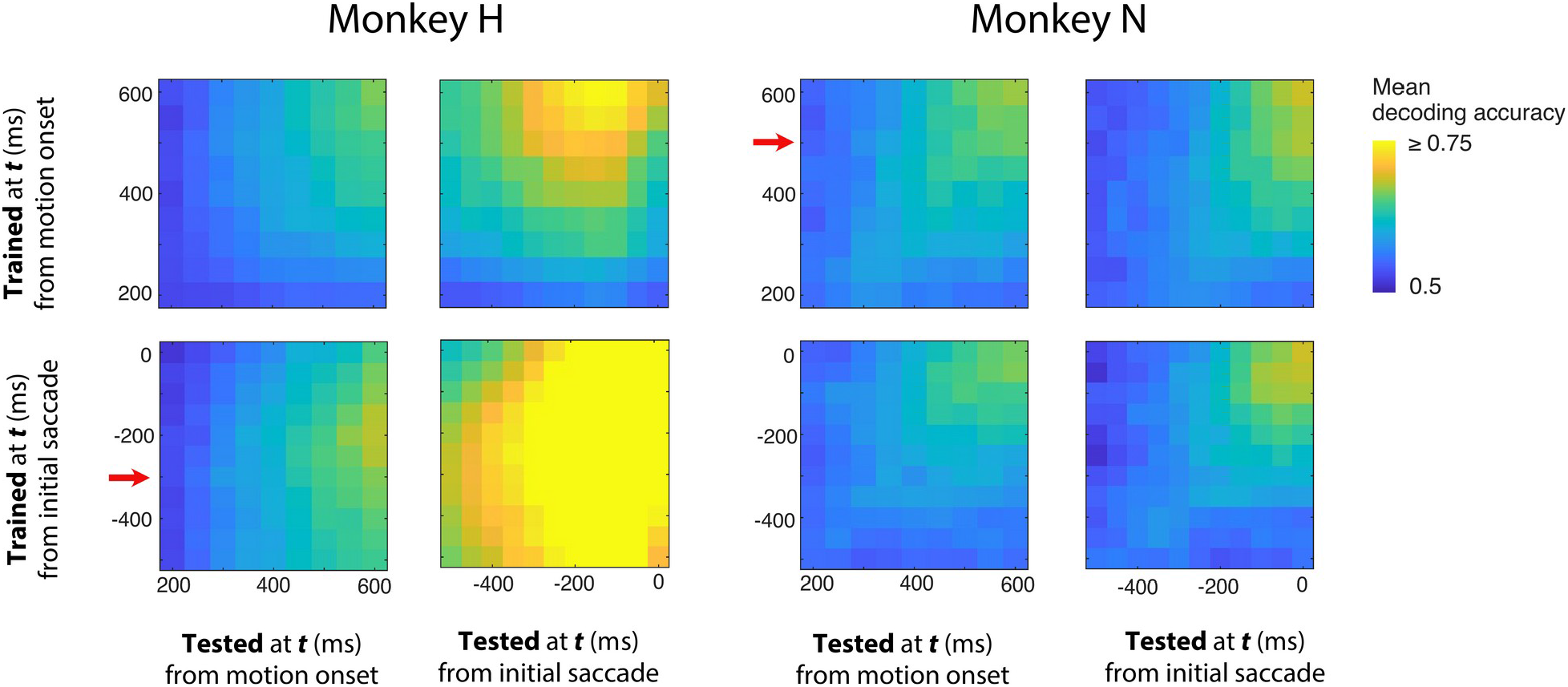
Temporal generalization of the What coding direction. Temporal generalization of choice decoders trained at different times during decision formation for monkey H (*left two columns*) and monkey N (*right two columns*). Each decoder was trained to discriminate the monkey’s eventual choice using population activity within a 100-ms window and evaluated on held-out trials across time. The *top row* shows decoders trained using activity aligned to motion onset and tested on population activity aligned either to motion onset (*left*) or to the *T*^when^ saccade (*right*). The *bottom row* shows the corresponding analyses for decoders trained using activity aligned to the *T*^when^ saccade. Thus, each heatmap depicts the generalization of *What* coding directions defined at different training times (vertical axis) to population activity sampled at different test times (horizontal axis). Each pixel shows the decoding accuracy averaged across recording sessions, with sessions weighted equally; chance performance is 0.5. The *What* CD used for subsequent analyses was selected to provide broad temporal generalization across the decision epoch while favoring temporal separation from the *T*^when^ saccade, thereby reducing potential contributions from peri-saccadic activity. Red arrows indicate the selected training windows for each monkey (monkey H: 300ms before the *T*^when^ saccade; monkey N: 500ms after motion onset).

## Footnotes

1 This prediction assumes that only the reporting structure changes—not the evidence or the computation that evaluates it.

2 A related phenomenon has been described in monkeys, where population decoding of prefrontal activity identified apparent sign reversals in the decoded decision variable prior to the final saccadic report [25]. Unlike the continuously monitored hand trajectory available in human subjects [24], however, this evidence for a change of mind in the monkey is inferred from population decoding rather than observed directly in an overt behavioral trace.

3 Some authors prefer “response time” to “reaction time,” reserving the latter for simple responses to stimulus onset and arguing that the former better describes tasks, like this one, in which the response reflects a decision process rather than a mere reaction [41]. We use “reaction time” throughout for consistency with common usage in the decision-making literature.

